# Recent Y chromosome divergence despite ancient origin of dioecy in poplars (*Populus*)

**DOI:** 10.1101/011817

**Authors:** Armando Geraldes, Charles A. Hefer, Arnaud Capron, Natalia Kolosova, Felix Martinez-Nuñez, Raju Y. Soolanayakanahally, Brian Stanton, Robert D. Guy, Shawn D. Mansfield, Carl J. Douglas, Quentin C. B. Cronk

**Author notes:** Equal contribution (Armando Geraldes, Charles A. Hefer). Corresponding author: Armando Geraldes, 6270 University Boulevard – Botany Department, UBC Vancouver, BC V6T 1Z4 Canada email: geraldes_at_mail_dot_ubc_dot_ca. Supplementary information: All supplementary files will be made available upon request. Data Accessibility: All sequence data will be deposited on Genbank and SRA and all other data files on Dryad.

## Abstract

All species of the genus *Populus* (poplar, aspen) are dioecious, suggesting an ancient origin of this trait. Theory suggests that non-recombining sex-linked regions should quickly spread, eventually becoming heteromorphic chromosomes. In contrast, we show using whole genome scans that the sex-associated region in *P. trichocarpa* is small and much younger than the age of the genus. This indicates that sex-determination is highly labile in poplar, consistent with recent evidence of “turnover” of sex determination regions in animals. We performed whole genome resequencing of 52 *Populus trichocarpa* (black cottonwood) and 34 *P. balsamifera* (balsam poplar) individuals of known sex. Genome-wide association studies (GWAS) in these unstructured populations identified 650 SNPs significantly associated with sex. We estimate the size of the sex-linked region to be ~100 Kbp. All significant SNPs were in strong linkage disequilibrium despite the fact that they were mapped to six different chromosomes (plus 3 unmapped scaffolds) in version 2.2 of the reference genome. We show that this is likely due to genome misassembly. The segregation pattern of sex associated SNPs revealed this to be an XY sex determining system. Estimated divergence times of X and Y haplotype sequences (6-7 MYA) are much more recent than the divergence of *P. trichocarpa* (poplar) and *P. tremuloides* (aspen). Consistent with this, in *P. tremuloides* we found no XY haplotype divergence within the *P. trichocarpa* sex-determining region. These two species therefore have a different genomic architecture of sex, suggestive of at least one turnover event in the recent past.

## Introduction

The separation of male and female sexual function into different individuals (dioecy) is an efficient way to ensure that sexual reproduction results in the recombination of genetic information from different individuals and is common in eukaryotes, occurring in 94% of animals [1] but only in about 6% of flowering plant species [1, 2]. Dioecy usually evolves from a cosexual ancestral state and involves at least two mutations. In one model, the pathway to XY systems involves one recessive mutation that suppresses male function (M^F^ -> M^s^) and a dominant mutation that suppresses female function (F^f^-> F^S^) [3], where the Y chromosome harbors the alleles M^F^ and F^S^ and the X chromosome the alleles M^s^ and F^f^. Recombination suppression between these loci on the Y chromosome likely evolves under the action of natural selection because recombination generates unfit sterile individuals [4]. With time, recombination suppression may extend to the rest of the chromosome via the accumulation of sexually antagonistic mutations on the Y [5], leading to the degeneration of the heterogametic sex chromosome (the Y or the W) via Muller’s ratchet, background selection and hitchhiking [6]. Under this view, old sex chromosomes are structurally and genetically divergent. The mammalian Y chromosome, having evolved ~170 MYA [7], is one such case of a degenerate Y chromosome that retains only a small fraction of the genes thought to be present in the autosomal pair from which the Y arose [8].

Studying old and degenerate Y chromosomes allows only for retrospective insights into their evolutionary origins. In some groups, sex chromosomes may be young and therefore provide windows into the initial stages of their evolution (e.g., [9, 10]). In plants, dioecy evolved independently in several clades allowing for a comparative approach that may reveal commonalities and peculiarities among independent origins of sex chromosomes [11]. Despite recent progress in the use of genomic resources to unravel the genetic basis of dioecy in plants such as papaya and white campion, the nature of sex-determining regions and sex-determining genes in plants remains elusive [12].

*Populus* species (poplars, cottonwoods and aspens) present an excellent opportunity to study the evolution of sex chromosomes. *Populus* and *Salix*, sister genera in the Salicaceae, are composed exclusively of dioecious species (with reports of rare cosexual genotypes, e.g. [13]), consistent with a single ancient origin of dioecy in this group around 65 MYA [14]. The cytological evidence (reviewed in [15]) for the existence of heteromorphic sex chromosomes is mixed, but in general there is no strong evidence for their existence (or for different chromosome counts in males and females), and the nature of the sex-determining region in *Populus* has remained elusive. Previous genetic mapping studies have mapped the sex-determining region to the proximal telomeric end of chromosome 19 in poplars and cottonwoods (*Populus* sections Tacamahaca and Aigeiros, [16, 17]) or to a pericentromeric region in aspens (*Populus* section Populus, [[18-20]). Some studies have proposed that females are the heterogametic sex (ZW system, [16, 19]) while other evidence suggests that males are (XY system, [17, 18, 20–22]). Recently, markers associated with sex were described for aspens, corresponding to the presence of the gene *TOZ19* on the Y chromosome of *P. tremula* and *P. tremuloides* and its absence from the X chromosome [22]. Here we use a genome-wide association approach (GWAS) to determine the genomic architecture of sex in two species of poplar.

## Results

### Genome-wide association analysis (GWAS)

We performed a simple case control GWAS between allele frequency at 3,656,736 loci with MAF>0.1 (minor allele frequency) and GR>0.9 (genotyping rate, Table S1) and sex (male vs. female) of 34 female and 18 male *P. trichocarpa* individuals (hereafter T52 association population, SI). After Bonferroni correction we recovered 623 single nucleotide polymorphisms (SNPs) significantly associated with sex (α<0.05; Fig. 1 and Table S2). Across all significant SNPs and accessions, females were homozygous at 99.9% of the genotypes and males were heterozygous at 94.0% of the genotypes, a pattern consistent with an XY sex determining system (Table 1).

**Fig. 1.**
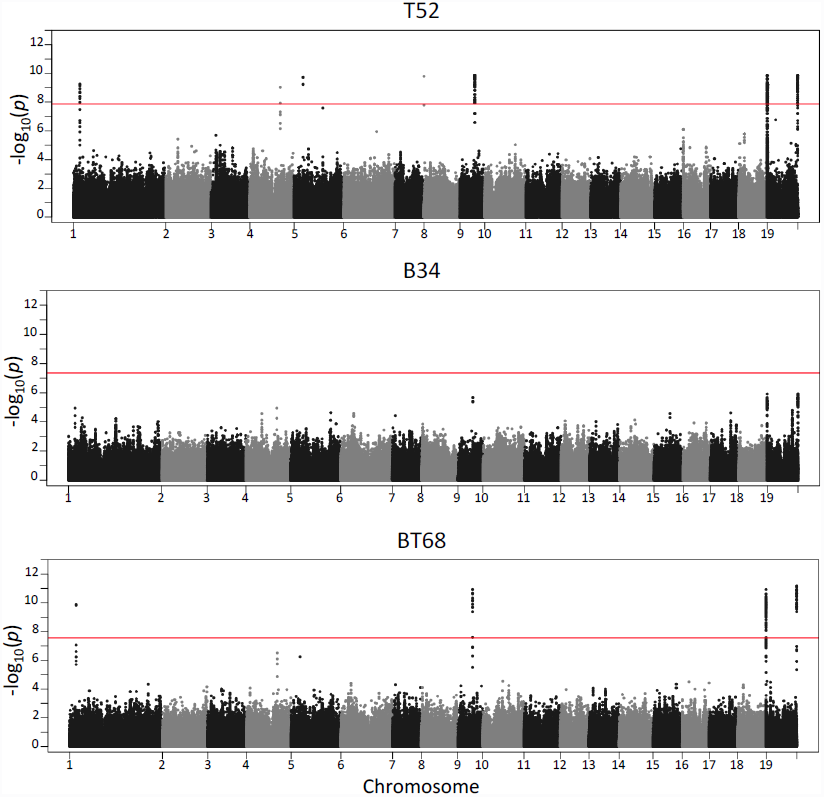

Manhattan plots depicting the GWAS results for association between allele frequency (v2.2 of the reference genome) and sex in three populations: 34 female and 18 male *P. trichocarpa* accessions (T52), 18 female and 16 male *P. balsamifera* accessions (B34) and 36 female and 32 male accessions, where half the samples of each sex are *P. trichocarpa* and the other half are *P. balsamifera* (BT68). SNPs mapped to unassembled scaffolds are not represented. The horizontal line indicates the –log_10_ (*p* value) corresponding to α<0.05 after Bonferroni correction for multiple testing.

**Table 1.**
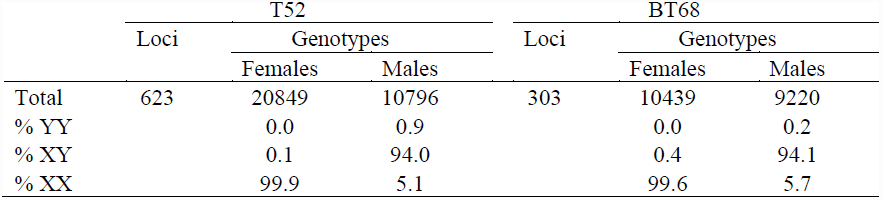
Number of loci associated with sex (and percent observed genotypes) in T52 and BT68.

|  | T52 |  |  | BT68 |  |  |
| --- | --- | --- | --- | --- | --- | --- |
|  | Loci | Genotypes |  | Loci | Genotypes |  |
|  |  | Females | Males |  | Females | Males |
| Total | 623 | 20849 | 10796 | 303 | 10439 | 9220 |
| % YY |  | 0.0 | 0.9 |  | 0.0 | 0.2 |
| % XY |  | 0.1 | 94.0 |  | 0.4 | 94.1 |
| % XX |  | 99.9 | 5.1 |  | 99.6 | 5.7 |

A similar analysis for 1,140,437 SNPs (Table S1) and sex (18 female and 16 male individuals, SI) in *P. balsamifera* (hereafter B34) recovered no SNPs statistically associated with sex (Fig. 1 and Table S2). Inspection of the results of the two analyses revealed that for 72.6% (452/623) of the significantly associated SNPs in *P. trichocarpa*, no data was available in *P. balsamifera* (i.e. SNPs had GR<0.9 and/or MAF<0.1). For the remaining SNPs, the vast majority (157/171) showed a similar pattern to that of SNPs significantly associated with sex in *P. trichocarpa*, i.e. females were homozygous and males heterozygous (with less than 10% of accessions deviating from this pattern) and the observed uncorrected p-values range was 2.07x10^-4^ -1.22x10^-6^ (Fig. 1 and Table S2).

Finally, we created a third association population consisting of 36 females and 32 males where, in each sex, equal numbers of accessions were *P. trichocarpa* and *P. balsamifera* (hereafter BT68, SI). In this population there were 1,782,995 SNPs with MAF>0.1 and GR>0.9 (Table S1) and 303 SNPs were significantly associated with sex (α<0.05; Fig. 1 and Table S2), of which only 27 were not significant in the analysis with *P. trichocarpa* alone (T52, Table S2). Across all significant SNPs and accessions in BT68, females were homozygous at 99.6% of the genotypes and males were heterozygous at 94.1% of the genotypes, a pattern again consistent with an XY sex determining system (Table 1).

In all three cases, Q-Q plots (Fig. S1) did not reveal an inflation of observed p-values with regards to the expected distribution of p-values, except for the extreme observed p-values. This is as expected given that T52 and B34 are unstructured populations and the population structure observed in BT68 did not co-vary with the phenotype (SI).

### Genomic distribution of sex-associated SNPs

Surprisingly, SNPs significantly associated with sex were located in 10 different regions of v2.2 of the *P. trichocarpa* reference genome assembly. The majority of SNPs associated with sex in T52 were located in the proximal end of chromosome 19 (hereafter Chr19P, 62.12%, 387/623) and the distal end of the same chromosome (hereafter Chr19D, 14.60%, 91/623). Remaining SNPs were located on chromosomes 1, 4, 5, 8, 9 and scaffolds 261, 1817 and 2325 (Fig. 1 and Table 2). Despite being distributed across different genomic regions, pairwise estimates of linkage disequilibrium between SNPs associated with sex were very high (all significant SNPs, average r^2^ =0.93, range 0.46-1; significant SNPs in putatively different genomic regions, average r^2^ =0.90, range 0.46-1; significant SNPs in the same genomic region, average r^2^ =0.97, range 0.51-1; Fig. S2).

**Table 2.**
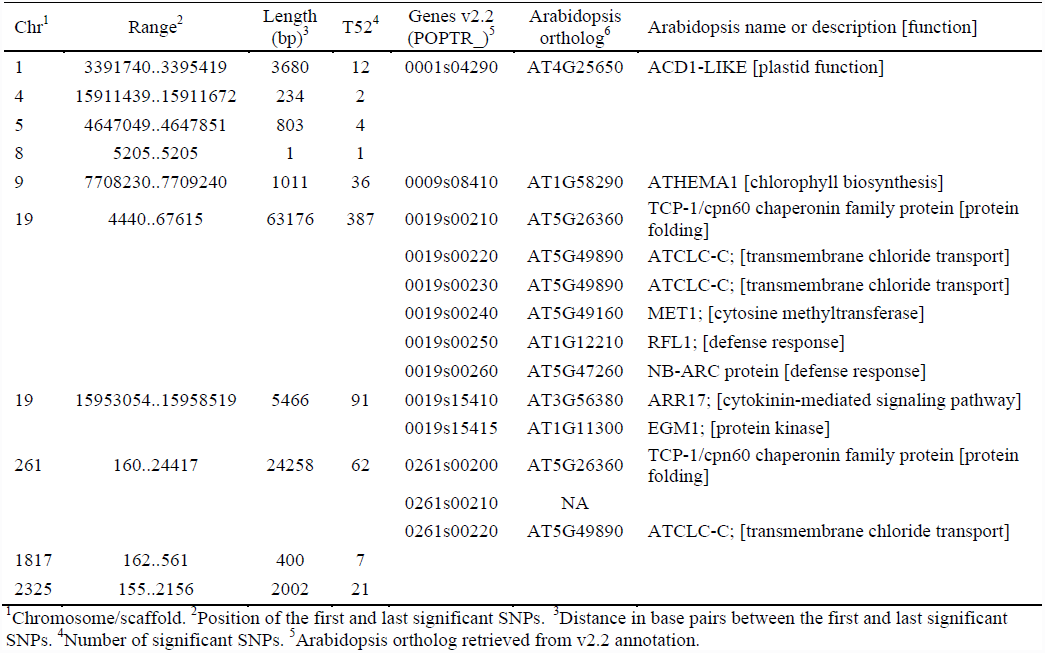
All regions significantly associated with sex in T52 (v2.2 of the genome annotation).

| Chr<sup>1</sup> | Range<sup>2</sup> | Length (bp)<sup>3</sup> | T52<sup>4</sup> | Genes v2.2 (POPTR_<sub>5</sub>)<sup>5</sup> | Arabidopsis ortholog<sup>6</sup> | Arabidopsis name or description [function] |
| --- | --- | --- | --- | --- | --- | --- |
| 1 | 3391740..3395419 | 3680 | 12 | 0001s04290 | AT4G25650 | ACD1-LIKE [plastid function] |
| 4 | 15911439..15911672 | 234 | 2 |  |  |  |
| 5 | 4647049..4647851 | 803 | 4 |  |  |  |
| 8 | 5205..5205 | 1 | 1 |  |  |  |
| 9 | 7708230..7709240 | 1011 | 36 | 0009s08410 | AT1G58290 | ATHEMA1 [chlorophyll biosynthesis] |
| 19 | 4440..67615 | 63176 | 387 | 0019s00210 | AT5G26360 | TCP-1/cpn60 chaperonin family protein [protein folding] |
|  |  |  |  | 0019s00220 | AT5G49890 | ATCLC-C; [transmembrane chloride transport] |
|  |  |  |  | 0019s00230 | AT5G49890 | ATCLC-C; [transmembrane chloride transport] |
|  |  |  |  | 0019s00240 | AT5G49160 | MET1; [cytosine methyltransferase] |
|  |  |  |  | 0019s00250 | AT1G12210 | RFL1; [defense response] |
|  |  |  |  | 0019s00260 | AT5G47260 | NB-ARC protein [defense response] |
| 19 | 15953054..15958519 | 5466 | 91 | 0019s15410 | AT3G56380 | ARR17; [cytokinin-mediated signaling pathway] |
|  |  |  |  | 0019s15415 | AT1G11300 | EGM1; [protein kinase] |
| 261 | 160..24417 | 24258 | 62 | 0261s00200 | AT5G26360 | TCP-1/cpn60 chaperonin family protein [protein folding] |
|  |  |  |  | 0261s00210 | NA |  |
|  |  |  |  | 0261s00220 | AT5G49890 | ATCLC-C; [transmembrane chloride transport] |
| 1817 | 162..561 | 400 | 7 |  |  |  |
| 2325 | 155..2156 | 2002 | 21 |  |  |  |

<sup>1</sup>Chromosome/scaffold. <sup>2</sup>Position of the first and last significant SNPs. <sup>3</sup>Distance in base pairs between the first and last significant SNPs. <sup>4</sup>Number of significant SNPs. <sup>5</sup>Arabidopsis ortholog retrieved from v2.2 annotation.

### Evidence for a single sex-specific locus in *P. trichocarpa*

Previous QTL experiments in *Populus* mapped sex to a single location suggesting inheritance as a single genetic locus [15]. Furthermore, sex always mapped to chromosome 19, albeit to different positions on the chromosome in different crosses/species [15]. Thus, the presence of SNPs associated with sex in different genomic regions in our GWAS might be due to problems with the assembly of the reference genome v2.2. To address this we repeated the GWAS for the T52 population after read mapping and SNP calling to genome assemblies v1.0 and v3.0. In both cases, sex again mapped to multiple regions although the details of the locations differed among assemblies (Fig. S3 and Tables S3-S4). The sex-determining region was therefore highly unstable with respect to assembly version.

We also performed cross mapping of sex-linked regions among assemblies with a BlastN (E-value cutoff 10^-10^, best 10 hits kept) search of the regions containing significant SNPs in v2.2 against assemblies v1.0 and v3.0 (Table S5). The resulting alignments indicated that the Chr09 and Chr19D sex-linked regions have similar locations in all three assemblies. All other sex-linked regions mapped to a different location in at least one of the three assemblies, e.g., the sex-linked region in Chr19P (v2.2 and v1.0) has moved to the distal end of Chr18 in v3.0.

Finally, we queried (BlastN, E–value cut–off 10^−6^, best hit kept) the BAC end sequences of a male *P. trichocarpa* library [23] to look for BAC clones nearby our sex-linked regions in which the two ends mapped to different locations in v2.2 and we identified 17 such BACs (SI). For three of these BACs, both ends were sex-linked. One BAC-end sequence from clone POR18-C06 maps to Chr19D and the other end maps to Chr01. One BAC-end sequence from clone POR02-A02 maps to Chr19P and the other end maps to scaffold 2325. Finally, one BAC-end sequence from clone POR07-E07 maps to Chr19P and the other end maps to Chr08. These results suggest that in this male, the sex-linked regions in Chr19D and Chr01, as well as in Chr08, Chr19P and scaffold 2325 are physically linked.

The above evidence, taken together, strongly suggests that assembly problems are sufficient to explain the genomic distribution of the sex-associated markers.

### Sex-linked regions in other accessions and species

We developed two PCR-RFLP assays for rapid genotyping of accessions in two of the regions with SNPs significantly associated with sex (Chr09 and Chr19P). Application of these assays to 8 samples of each sex and species used in the GWAS revealed full agreement between WGS and PCR-RFLP inferred genotypes (Fig. S4) and confirmed that one male of each species, BELA18-5 and AP2446, appears to be recombinant; i.e. both of these males are homozygous for the majority of SNPs in Chr19P, but are heterozygous for significant SNPs in the other sex-linked regions (Fig. S2). Application of these assays to *P. trichocarpa* and *P. balsamifera* accessions of each sex that were not used in the GWAS showed that these SNPs are linked with sex in independent accessions (Fig. S4). Finally, we used these assays to determine whether these SNPs are also linked to sex in other species. All 16 *P. deltoides* and 16 *P. nigra* accessions of known sex assayed were homozygous (XX) in females and heterozygous (XY) in males (Fig. S4). This indicates that the *P. trichocarpa/P. balsamifera* sex-linked markers are conserved in these species. However, for one female and three male *P. tremuloides* accessions no differences between sexes were observed (Fig. S4), suggesting that in aspens these regions are not sex-linked.

### Phylogeny of X and Y alleles

We performed allele-specific amplification and sequencing of X and Y alleles in two regions associated with sex (gene POPTR_0019s00240 on Chr19P and gene POPTR_0009s08410 on Chr09) using several males from each of four species: *P. trichocarpa*, *P. balsamifera*, *P. deltoides* and *P. nigra* (hereafter referred to as “cottonwoods”). We also included sequences cloned from *P. tremuloides* (hereafter referred to as “aspen”), the reference genome sequence, as well as the genome sequence of the paralog of each gene that resulted from the Salicoid whole genome duplication (WGD) event [24]. Maximum likelihood phylogenies of each region (Fig. 2 and Fig. S5) show that both X and Y chromosome alleles from all four cottonwood species group by gametolog (i.e., X or Y) and not by species, indicating that X and Y chromosome alleles began to diverge before species did. Note that because for one of the amplicons in Chr19P we failed to amplify the X gametolog of *P. nigra, P. nigra* alleles are not shown in the concatenated phylogeny of Chr19P (Fig. 2); nevertheless phylogenies of the other two amplicons in Chr19P show unequivocally that *P. nigra* alleles cluster by gametolog (Fig. S5). The placement of aspen alleles with respect to X and Y alleles from cottonwoods is uncertain. For the region in Chr09, they cluster with cottonwood sequences from the X gametolog, but with low bootstrap support, while for the region in Chr19P they appear basal to the X and Y clades (Fig. 2).

**Fig. 2.**
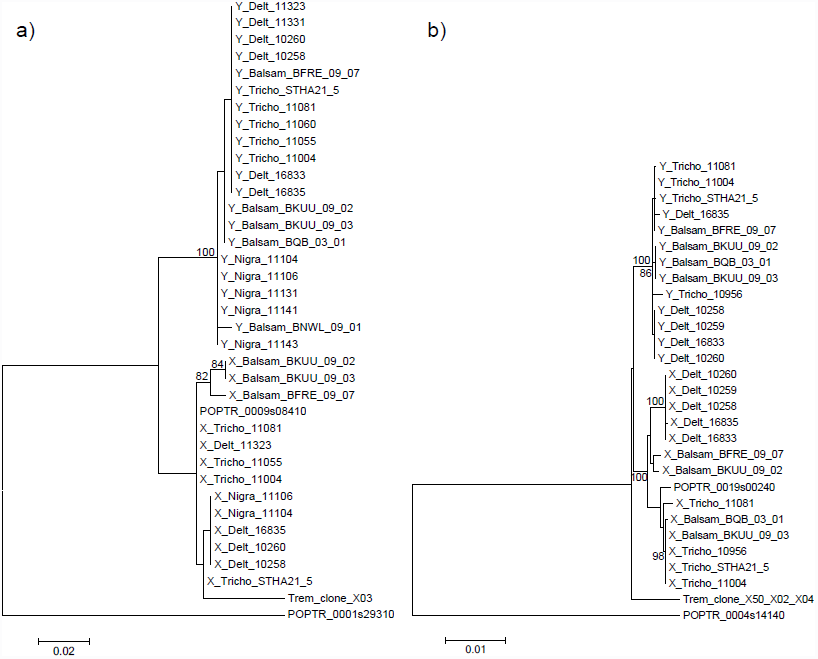
Neighbor joining maximum likelihood phylogenies of regions significantly associated with sex. For each phylogeny, only male accessions were used. Y chromosome alleles are indicated with a Y and X chromosome alleles with an X at the beginning of the sequence name, followed by species and accession identifiers. Only one random *P. tremuloides* allele is depicted. Phylogenies including all *P. tremuloides* sequences are available in Fig. S5. Each phylogeny also includes the *P. trichocarpa* reference sequence from genome assembly v2.2 (POPTR_0009s08410 and POPTR_00019s00240) and the reference sequence from genome assembly v2.2 of the paralog from the Salicoid WGD (POPTR_0001s29310 and POPTR_0004s14140). Only bootstrap values higher than 80 are shown. a) Phylogeny of Chr09 region (Amplicon1:Chr09:7690067, SI) and b) Concatenated phylogeny of Chr19 (Amplicon1:Chr19:40024, Amplicon2:Chr19:41515 and Amplicon3:Chr19:44107, SI).

### Divergence at X and Y regions

The phylogenies in Fig. 2 clearly suggest that recombination between the X and Y regions identified here ceased, and their divergence in cottonwoods started, after the split between cottonwoods and aspens. The amount of divergence at silent sites (K_s_), between the X and Y clade (Chr09 K_s_ =0.0224 and Chr19P K_s_ =0.0163) was only slightly lower than K_S_ between all XY cottonwood alleles and aspen (Chr09 K_s_ =0.0638 and Chr19P K_s_ =0.0186), and both were roughly one tenth the K_s_ between the XY clade and the paralog from the Salicoid WGD (Chr09 K_s_ =0.2027 and Chr19P K_s_ =0.1774; Table 3). Assuming the timing of the WGD to be 65 MYA [24], then XY divergence for Chr9 would be approx. 7.2 MYA and Chr19 divergence approx. 6.0 MYA.

**Table 3.** Divergence estimates at two regions associated with sex.

|  |  | X-Y<sup>1</sup> | XY<sup>2</sup> |  | X<sup>3</sup> |  | Y<sup>4</sup> |  |
| --- | --- | --- | --- | --- | --- | --- | --- | --- |
|  |  |  | *P.*<br>*tremuloides* | Salicoid<br>Paralog | *P.*<br>*tremuloides* | Salicoid<br>Paralog | *P.*<br>*tremuloides* | Salicoid<br>Paralog |
| Chr09 | K<sub>s</sub><sup>5</sup> | 0.0224 | 0.0638 | 0.2027 | 0.0547 | 0.2174 | 0.0683 | 0.1889 |
|  | K<sub>a</sub>/K<sub>s</sub><sup>6</sup> | 1.606 | 0.761 | 0.667 | 0.56 | 0.582 | 0.87 | 0.737 |
| Chr19 | K<sub>s</sub><sup>5</sup> | 0.0163 | 0.0186 | 0.1774 | 0.0227 | 0.182 | 0.0144 | 0.1728 |
|  | K<sub>a</sub>/K<sub>s</sub><sup>6</sup> | 0.295 | 0.331 | 0.198 | 0.247 | 0.188 | 0.463 | 0.208 |

<sup>1</sup>Divergence between the X and Y clades from Fig. 2. <sup>2</sup>Divergence between all cottonwood sequences and *P. tremuloides*/Salicoid paralog. <sup>3</sup>Divergence between all sequences from the X lineage in cottonwoods and *P. tremuloides*/Salicoid paralog. <sup>4</sup>Divergence between all sequences from the Y lineage in cottonwoods and *P. tremuloides*/Salicoid paralog. <sup>5</sup>Synonymous substitutions at synonymous sites included in the estimation of K. <sup>6</sup>The ratio of non-synonymous substitutions per non-synonymous site to synonymous substitutions per synonymous site.

For both regions, the ratio of non-synonymous substitutions per non-synonymous site to synonymous substitutions per synonymous site (K_a_/K_s_) is higher for the Y lineage than for the X lineage (Table 3). This pattern is consistent with an accumulation of deleterious mutations following recombination suppression. The fact that this difference is larger when divergence is measured to aspen (Chr09 X K_a_/K_s_ =0.560, Y K_a_/K_s_ =0.870 and Chr19P X K_a_/K_s_ =0.247 and Y K_a_/K_s_ =0.463), than when divergence is measured to the Salicoid paralog (Chr09 X K_a_/K_s_ =0.582, Y K_a_/K_s_ =0.737 and Chr19P X K_a_/K_s_ =0.188 and Y K_a_/K_s_ =0.208), suggests that the increase in K_a_/K_s_ in the Y lineage is recent. Furthermore, despite its recent origin our data suggest that the Y-haplotype is already becoming non-functional as we observe frame-shift insertions/deletions in Y sequences of POPTR_0009s08410.

### Size and composition of the sex-linked region

The 650 sex-associated SNPs, if concatenated, cover a total genomic region of ~100 Kbp. Thus given the evidence above that a single region is involved, that region is extremely small. We considered if there might be large missing tracts of Y sequence that were not detected by our read-mapping protocol. *De novo* assembly of unmapped reads from male accessions revealed four male-specific contigs that are candidates for such Y sequences (SI). However these are short (longest contig is 2514 bp) and BlastN searches reveal that they either are repetitive in nature or have significant similarity to the sex-linked regions identified by GWAS. There is thus no present evidence that the sex-locus in *P. trichocarpa* is significantly larger than reported here. The 13 genes in the sex-linked region (Table 2) cover a range of functional classes, including DNA methylation, hormone regulation, ion transport and plant defense.

## Discussion

### XY sex-determining system

The identification of 650 sex-specific SNPs heterozygous in males and homozygous in females by GWAS unequivocally shows that an XY system is involved in sex-determination in *P. trichocarpa/P. balsamifera*. The findings were fully and independently supported by PCR/RFLP-assays for two representative SNPs that distinguish X and Y alleles carried out on *P. trichocarpa*, *P. balsamifera*, *P. deltoides*, and *P. nigra* individuals of known sex not included in the GWAS. This finding of an XY system in cottonwoods (*Populus* sections Tacamahaca and Aigeiros) is further supported by previous reports of an XY system in *P. nigra* of section Aigeiros [17] and aspens of section Populus [20–22] but is at odds with previous suggestions that a ZW (female heterogamy) system of sex determination may function in *P. trichocarpa* [16].

The previous suggestion that *P. trichocarpa* has a ZW system was based on inferences from a cross of *P. deltoides* x (*P. nigra* x *P. deltoides*) and was not supported by sex-specific markers [16]. Our results run counter to those inferences, but it is conceivable that a ZW system, with a highly divergent W chromosome that is not represented in the *P. trichocarpa* reference sequence [24], could produce the observed pattern of homozygosity in females and heterozygosity in males at SNPs significantly associated with sex, as the W sequence would be absent in males and divergent enough from the Z that reads from the W chromosome do not map to the reference sequence. Thus, in females, apparent homozygosity would in fact be due to hemizygosity. Several observations contradict this hypothesis: a) we observed heterozygous positions in females at sex-linked regions intermingled with SNPs significantly associated with sex (SI), b) Sanger sequencing of females for sex-linked regions revealed heterozygous positions (SI), c) qPCR of two sex-linked regions (Chr19P and Chr19D) revealed a 1:1 ratio of amplification of autosomal to sex-linked regions in both sexes (SI), d) WGS coverage is approximately similar in males and females at sex-linked regions (SI) and e) *de novo* assembly of female specific regions did not reveal unassembled regions unique to females (SI). Given strong direct evidence for an XY system from sex-linked markers, and absence of evidence for hemizygosity in females, we now argue that the ZW hypothesis can be discounted.

### Genomic architecture of the sex locus

The 623 sex-specific SNP markers identified by GWAS in T52 are in nearly complete genetic linkage (Fig. S2). The majority of these markers map to Chr19P confirming previous studies that implicate this region as the location of the sex locus [16, 17]. However, remarkably, we found that sex-linked markers in apparent genetic linkage map to multiple physical locations in the three *P. trichocarpa* genome assemblies (Fig. 1 and Tables S2-S4). Our data do not support the existence of a multi-locus system of sex determination in *P. trichocarpa*, but instead suggest that a single genetic region controls dioecy and that the genome assembly is a work in progress with some contigs from Chr19P having been misassembled into other genomic regions. Sex determining regions and sex chromosomes are notoriously difficult to assemble [25]. Further refinement of the assembly regarding the sex locus may require complementary methods.

### The age of the cottonwood sex locus and evolution of dioecy in *Populus*

We find the same sex-linked markers in *P. trichocarpa* and *P. balsamifera* (*Populus* section Tacamahaca) as well as in *P. nigra* and *P. deltoides* (*Populus* section Aigeiros). The sex locus therefore predates the divergence of these species. Sequence analysis of sex regions of these species suggests an approximate date of 6-7 MYA (late Tertiary) for the divergence of X and Y. The fossil record indicates that aspens (*Populus* sect. Populus) likely diverged from cottonwoods long before this, given that middle to late Oligocene (~25 MYA) fossils of section Populus from Alaska have been reported [26]. Consistent with this is the fact that the polymorphic loci we identified do not provide sex-specific markers in aspens. Furthermore, sex linked markers have recently been identified in the pericentromeric region of Chr19 in aspens [22]. Genes in this region are not homologous to sex-linked genes identified in our study, and SNPs in this region do not segregate with sex in our mapping populations (SI); hence aspens and cottonwoods likely have independent sex determining mechanisms.

If there were a single origin of dioecy in this group, it is problematic that there are apparently distinct sex-determining loci in *Populus*. One plausible explanation is that there has been at least one sex-determination mechanism “turnover” since the divergence of poplars and aspens. The labile nature of sex determining regions is well known, with many examples of "turnover" of sex determining regions from diverse groups [27]. Mapping of sex-linked regions in other *Populus* species as well as in the sister genus *Salix* (willows) would provide further insight into the dynamics of sex-linked region turnover in the Salicaceae.

### The size of the sex locus

One remarkable feature of the sex locus described here is its compactness. Concatenating all the regions with sex specific markers leads to a total estimated size for the sex-determining region of ~100 Kbp. This small size is consistent with the difficulties encountered in finding sex-specific markers in the Salicaceae (reviewed in [15]). However, there are good reasons for supposing that a non-recombining region at a sex locus will rapidly expand, eventually to encompass an entire chromosome [28]. Such expansion is empirically well documented in other plant systems [29] and is driven by sexual conflict making it advantageous for more and more genes to be captured by the non-recombining regions. Even the 6-7 MYA date we estimate for the divergence of X and Y alleles would likely be sufficient for expansion to encompass a considerable portion of a chromosome. Therefore the apparent remarkably small size of the *P. trichocarpa* sex locus requires explanation.

One possibility is that the actual size of the cottonwood sex-determining locus is larger than it appears due to large tandem duplications and transposable element insertions in the Y. Yet, our *de novo* assembly of male-specific unmapped reads revealed only four small male-specific contigs (average length 1877 bp, SI) and these have either Blast hits to the sex-linked regions identified with GWAS (SI) or consist mostly of low complexity repetitive sequence. We were unable to retrieve further male-specific contigs, specifically, regions of higher divergence to the female reference sequence that may be indicative of older divergence strata as observed in other animal [8] and plant species [29]. Future investigations might reveal larger Y-specific regions. Alternatively, it is possible that features unique to trees dampen the expansion of sex determining regions. For instance, sexual conflict may be minimal in trees as carbon investment in reproduction is a relatively small annual cost compared to the massive storage of carbon in wood, a tissue with no obvious secondary sexual characteristics.

### Functional insights into sex-determination in cottonwoods

The sex-linked specific region in *P. trichocarpa* contains 13 genes (Table 2). However it is too early to say which, if any, of these genes are the master-regulators of sex. The reference genome is from a female (XX) individual [24] and, as suggested above, further work is required to fully characterize the Y chromosome. Furthermore, many of the genes in this region have poorly defined functions. Nevertheless, there are at least two plausible candidate genes. One, a poplar ortholog of the *Arabidopsis thaliana* [Arabidopsis] cytokinin pathway-associated *ARABIDOPSIS RESPONSE REGULATOR 17* (*ARR17*), is implicated in phytohormone signaling and the other, the poplar ortholog of Arabidopsis *METHYLTRANSFERASE 1*, (*MET1*), is involved in DNA methylation.

Phytohormone signaling is involved in other plant sex determination systems, such as the ethylene pathway in cucumber [30], and it is possible that cytokinin signaling, mediated by *ARR17* is used in poplar. DNA methylation has been implicated in sex determination in other plant systems, e.g. *Silene latifolia* [31]. In an andromonoecious clone of *P. tomentosa* expression of the poplar orthologue of *MET1* was significantly higher in all stages of female flower development [32]. In Arabidopsis, *MET1* is required for maintenance of epigenetic memory [33] and is involved in reproductive development including the control of floral homeotic genes such as *AGAMOUS*, *APETALA3* and *SUPERMAN* [34, 35].

Due to the Salicoid WGD [24] there are two paralogs of genes in the sex locus region such as *ARR17* and *MET1*, relative to Arabidopsis. Neofunctionalization, in which one copy has evolved a specific sex-determining function while the other copy retains the ancestral function, is therefore possible. The WGD may thus be important in the evolution of dioecy in this group. Functional differences between the paralog in the sex-specific region and an autosomal sister paralog could reveal pathways involved in sex determination.

## Materials and Methods

Tree sex was determined by visual inspection of flowers. DNA from *Populus trichocarpa* and *P. balsamifera* association populations was extracted from leaves and sequenced (100bp paired-end reads) on an Illumina HiSeq at the Genome Sciences Centre, Vancouver, BC to either 15x or 30x coverage (SI). Sequence data generated ranged from 31-241 million reads. All sequences are deposited at the NCBI short read archive under SRA XXX. Illumina reads were aligned to reference *P. trichocarpa* genome assemblies v1.0, v2.2 and v3.0 (http://www.phytozome.net) using BWA version 0.6.1 [36] with a 4 bp misalignment threshold, disallowing insertions or deletions within 5bp of the end of the sequence (aln –n 0.04 -i 5), maximum insert size of 500 bp (sampe -a 500), and default values for the remaining parameters. Paired-end mate information was synced using Picard-tools FixMateInformation (http://picard.sourceforge.net/). Local re-alignment was performed on identified regions with high SNP entropy, using a window-size of 10 bp, and a mismatch fraction of 0.15 for base qualities to identify mismatched regions using GATK version 1.5 [36]. Indel re-alignments were restricted to regions with a maximum insert size of 3 Kbp, and the maximum positional change of an indel set to 200 bp. Variant calls were made using the duplicate-marked alignment files and the UnifiedGenotyper from GATK emitting variant with a minimum phred-scaled confidence threshold of 30. We used vcftools [37] to filter out any variants where coverage was <5X and where more than two bases were segregating.

We performed a standard case/control GWAS between allele frequencies and sex phenotype using Plink v1.07 [38]. We report associations at α<0.05 after Bonferroni correction for multiple testing. Analysis of population structure in the three association populations is given in SI.

PCR-RFLP (polymerase chain reaction followed by restriction fragment length polymorphism) genotyping assays in two regions associated with sex were developed as follows: mpileup files were converted into fasta files by generating calls at each base of the reference whenever coverage at the position in each individual was higher than six and whenever heterozygote genotypes were present by requiring that each allele had coverage of at least three. All other positions were considered missing data. The fasta sequences were used to design PCR primers in regions conserved across all accessions to amplify two short fragments on the sex-linked regions that mapped to Chr09 and Chr19P. PCR primers, amplicons and protocol details are in SI. The Chr09 amplicon was digested with *Bsl*I (New England Biolabs, Ipswich, MA) and *Cla*I (New England Biolabs, Ipswich, MA); the Chr19 amplicon was digested with *TspR*I (New England Biolabs, Ipswich, MA); see SI for details. The same assays were used in *P. deltoides, P. nigra* and *P. tremuloides* accessions (SI).

To generate haplotypic Sanger sequences from selected male accessions (SI), allele-specific primers [39] were designed for three regions of the gene POPTR_0019s00240 on Chr19P and for one region of the gene POPTR_0009s08410 on Chr09 (SI). Each allele-specific primer was used with the common primer to generate an allele-specific PCR fragment that was subsequently cloned and Sanger sequenced. PCR protocol, amplicon and cloning details are in SI. Chromatograms from Sanger sequencing were visually inspected, trimmed, and aligned with BioEdit [40]. Sequences were aligned to the closest *P. trichocarpa* paralog (resulting from the Salicoid WGD); for Chr09 the paralog is POPTR_0001s29310, and for Chr19 the paralog is POPTR_0004s14140) and neighbor joining maximum likelihood trees for each amplicon were estimated in MEGA v5.03 [41] using the Tamura-Nei model and complete deletion of all sites with missing data and gaps. Levels of divergence were calculated for synonymous sites (K_s_) only and for replacement sites only (K_a_) in DNAsp v5 [42]. Sequence data is deposited in NCBI under accession numbers XXX.

## Acknowledgements

This work was supported by the Genome Canada Large-Scale Applied Research Program (POPCAN, project 168BIO) to QCBC, CJD, RDG and SDM. Additional support was received from a NSERC (Canada) Discovery Grant to QCBC. We thank Dr. Michael Friedmann for assistance with project management, Drs. Steffi Fritsche and Daisie Huang for assistance and discussion, Drs. Loren Rieseberg and Sally Otto for comments on the manuscript; Dr. Barbara Thomas for aspen material and both POPCAN project personnel, AAFC staff and Rich Shuren who kindly helped in recording tree sex.

